# The Impact of Remediation Through Stabilizing Amendments on Taxonomic and Metabolic Patterns of Bacteria and Archaea in Cadmium-Contaminated Paddy Fields in Southwestern China

**DOI:** 10.1101/311902

**Authors:** Yi Chen, LuLu Pan, QianQian Jiang, FangFang Chen, MengDi Xie, XiCong Lai, WenQing Chen, ZhiWei Tao

## Abstract

The in-situ immobilization of heavy metal contamination in soils using stabilizing amendments is a cost-effective remediation technique. However, previous research on mediating cadmium polluted fields using amendments has focused mainly on the absorption and accumulation of cadmium by plants, rather than the response of soil microorganisms to amendments. In our study, five compounds with different pH values and concentrations of carbonate minerals, clay minerals, sulfur, and phosphorous were selected to investigate their effect on the soil microorganisms and metabolic patterns through metagenomic analysis over three months in cadmium-contaminated paddy fields (CCPFs) in southwestern China. The results showed that the pH value of the amendments was the major factor determining the microbial diversity and communities. In weak acidic paddy fields, the appropriate use of an alkaline amendment composed mainly of calcium oxide increased the pH value of the soil, which helped to improve the soil microbial diversity, promote the growth of azotobacter, nitrate-reducing bacteria (such as *Pseudomonas),* and metabolisms of nitrogen fixation and reduction, which contributing to the decrease of available cadmium in soils. Acid amendments which significantly reduced the soil pH value, had lowest removal rate of available cadmium and showed significant restrictive effects on bacterial and arhaeal diversity and growth. In addition, the effects differed between different alkaline amendments. Alkaline amendment composed of magnesium oxide promoted the growth of *Pseudomonas*, but also inhibited *Nitrosospira* and the metabolism of soil nitrogen fixation. In conclusion, when applying amendments to remediate cadmium-contaminated paddy soils, we need to take into account the pH value of the amendment and the content of each component, and ensure the efficiency of amendments while at the same time maintaining a positive effect on soil microorganisms.

**Importance:** Previous research on amendment applications in cadmium-contaminated paddy fields (CCPFs) has mainly focused on the availability of cadmium to plants, the identification of the best functional additive proportion of immobilization remediation, and the absorption and accumulation of heavy metals by plant, etc. However, research on the taxonomic and metabolic patterns of bacteria and archaea in amended soils, especially purple paddy soil, which is typical of the Sichuan area (4.601 million hectares), is insufficient. Secondly, the plough layer of paddy soil would be in a reduction state during the period of irrigation, but be in an oxidation state when fields were in draining and drying periods. The periodical alternation of wetting and drying forms unique physical, chemical, and biological properties, and it is vital to understanding the taxonomic and functional dynamics of microbiomes in amended soils in order to improve the process performance.

## Introduction

Cadmium is characterized by its toxicity, accumulation, and mobility in the environment, and can result in potential hazards for human health through the soil-food chain transfer (1). In recent years, with the increases in atmospheric deposition and human activities, such as mining, the application of organic and inorganic fertilizers or sewage sludge, and wastewater irrigation, heavy metal pollution in soil has increased dramatically (2,3,4). In China, heavy metal pollution in soil is decreasing the country’s grain harvest by tens of millions of tons every year (5, 6). Another study showed that more than 70% of the rice grain samples from metal-contaminated rice fields across south China were Cd-contaminated (7). The multi-target regional geochemical survey of Chengdu economic zone revealed that the abundance of heavy metals, especially cadmium, was abnormal over large areas of Deyang (northeast of the Chengdu Plain). Besides, the results of the ecological geochemical assessment conducted in this region showed that the source of heavy metals pollution of soil, such as cadmium pollution, is the Longmen Mountains, which accumulates the mineral cadmium and near-surface atmospheric dust produced by industrial and mining enterprises. Heavy metal pollution has been a severe challenge to the sustainable utilization of cultivated land resource and food production safety. Therefore, environmental engineers have given high priority to the reduction of of toxic metal bioavailability in croplands to ensure food security and human health (8).

At present, the techniques of soil heavy metal pollution remediation are divided into four parts: phytoremediation, micro-remediation, physical remediation, and chemical remediation. The new-soil technology, adsorption, and electrokinetics (EK) belong to the physical remediation group, while leaching, biological reduction, complexation, and in-situ immobilization (9) are all part of the chemical remediation group. Of these remediation technologies, in-situ immobilization is considered to be the most effective and low-cost remediation, especially for non-point source pollution in cultivated soil (10). The key to the technology is to select suitable amendments, of which the most commonly used ones are alkaline materials, phosphorous materials, clay minerals, iron manganese oxides and organic materials (11). These materials can change the existing forms of heavy metal in soil through ion exchange, adsorption, and precipitation, and reduce the mobility and bioavailability of heavy metals. The environmental effect of soil amendment is an important problem; some amendment materials, such as phosphor, may cause environmental problems in certain environments. Zhao (12) found that the excessive use of phosphorus in farmland can produce some negative effects on soil microorganisms and induce zinc and calcium deficiency in crops, which could negatively affect the output of crops. Moreover, phosphorized materials may contain other heavy metals (such as calcium superphosphate), which can be easily introduced into the soil, and cause new soil heavy metal pollution. Previous research on Cd-contaminated paddy filed (CCPF) mainly focused on the impact of the applied amendments on the availability of cadmium to plants, identification of the best functional additive proportion of immobilization remediation, and absorption and accumulation of cadmium by plants; however, there have been limited studies on soil microorganisms and their metabolisms responding to amendments, especially in paddy fields rather than in the laboratory.

Therefore, we used a molecular community profiling tool (Illumina MiSeq system) and limited sequencing of selected clone libraries (V3-V4 regions of 16s rRNA) to evaluate soil bacterial and archaea communities. The aims of this study were to (i) investigate how bacteria and archaea composition and distribution vary between diverse amended soils; (ii) identify the active gene functions of microbial communities, especially the dominant functions with significant variation during remediation; and (iii) explore the relationship between the taxonomic and functional patterns of microbiomes from amended soils, especially functional bacterial, such as azotobacter and nitrate-reducing bacteria (NRB).

## Materials and Methods

### Experimental Fields and Amendments

The paddy soil selected for this study was typical purple paddy soil from Mianzhu, Sichuan Province (19.24°N, 105.71°E); with an area of 4.601 million hectares, the experimental fields account for 36.5% of the total area of cultivated land in Sichuan (13). The fields in this study were designated into two control fields (Fig. 1), one of which was left fallow without amendment (CK1) and the other planted with rice only (CK2), and five (A-E) experimental fields with areas of 9,375, 9,029, 7,665, 8,740, and 3,324 m^2^, respectively. The average pH value of those fields ranged from 6.12-6.57, indicating acidic soil. We remediated the CCPFs with five diverse amendments using materials (Table 1) that had been researched at least for three years in fields, but mostly in laboratories. They were acid amendment (E), neutral amendment (C) and three alkaline amendments of which A and B were mainly composed of CaO (calcium oxide) and D of MgO (magnesium oxide). The experiment comprised the entire process of rice planting from sowing in June 8 until September 25, when the rice was harvested. The mean annual precipitation was 74.86mm, with more than 60% falling from June-August, while the temperature during the experimental period ranged from 12°C-37°C (14).

**FIG 1.**
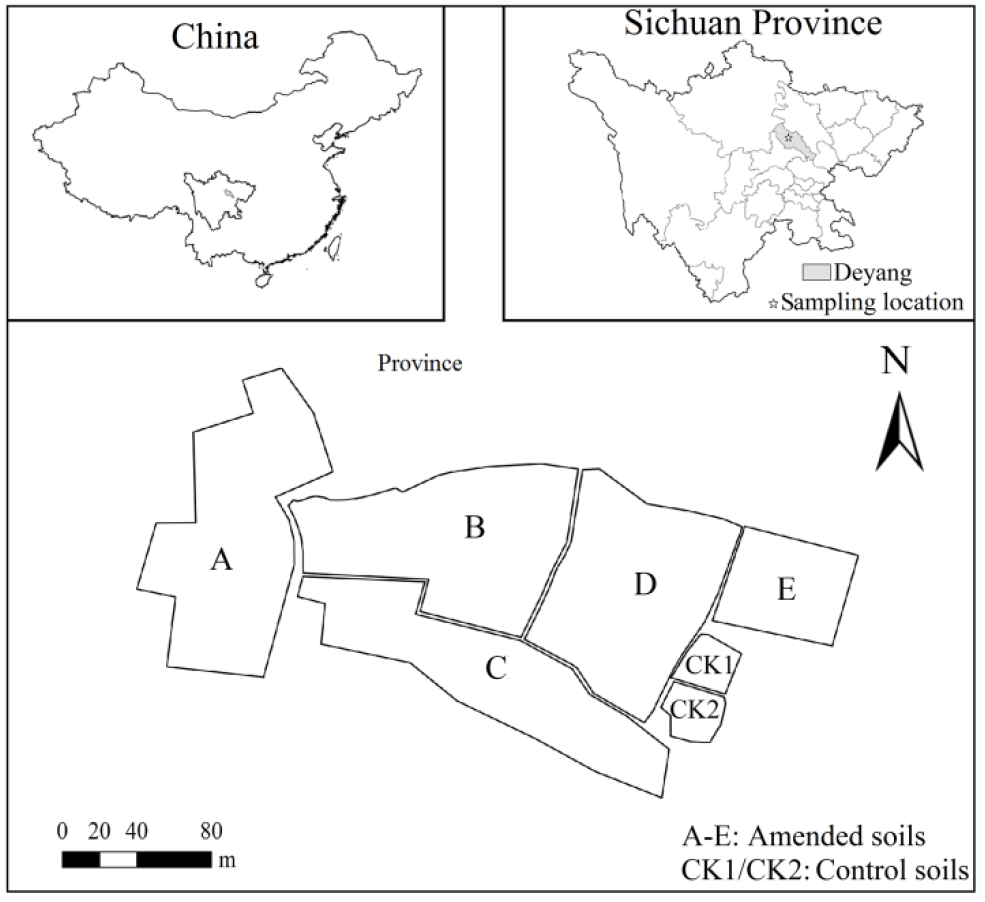
Location of amended soils in Sichuan, China.

**Table 1.**
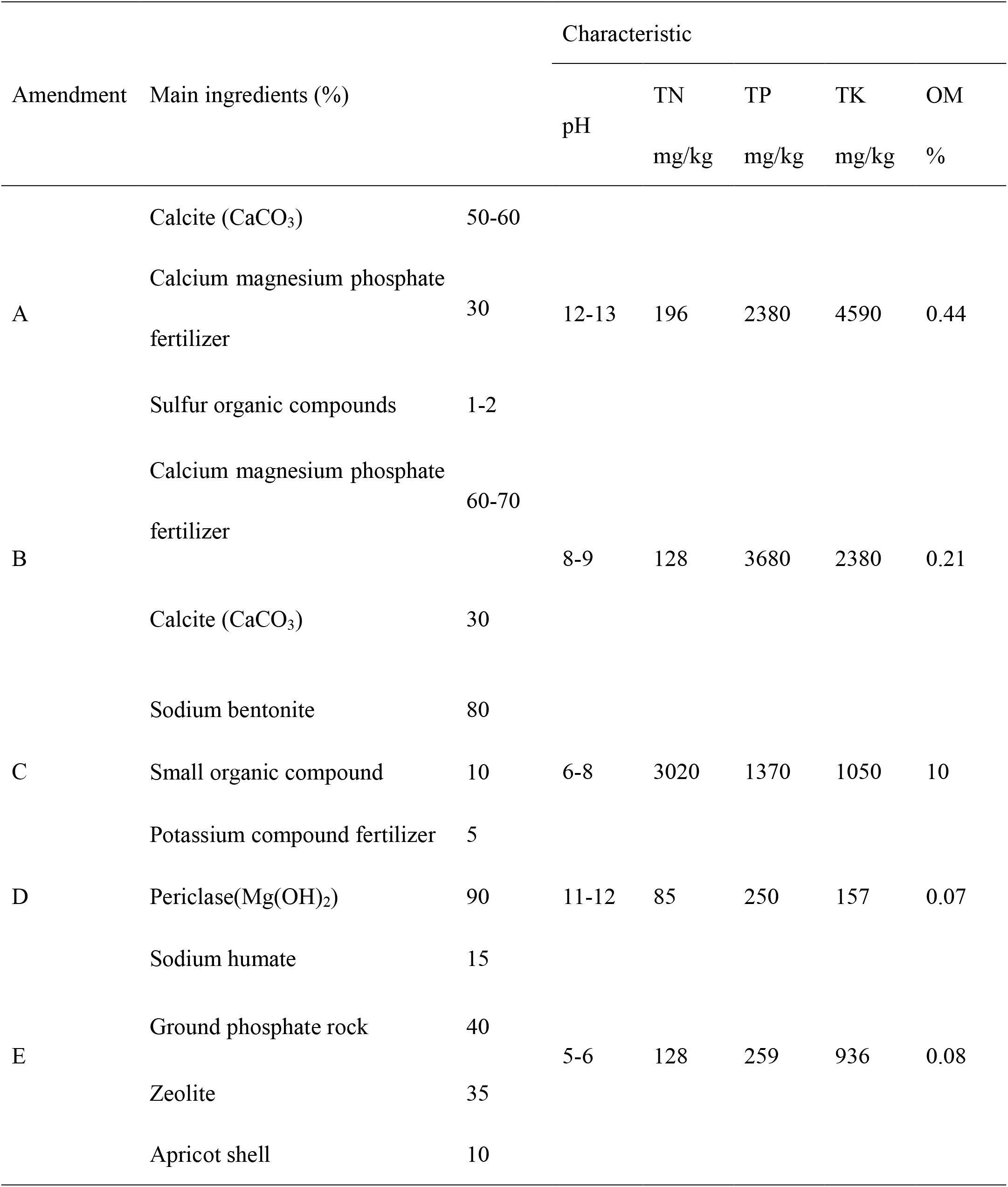
Main ingredientsand properties of the five amendments

### Sample Collection and Analyses

Five CCPFs were restored with five compound amendments and two were kept as controls. A total of 51, 58, 50, 60, 26, 7, and 7 samples were collected per site each time according to the grid method, with an average grid size of 10 m. The sampling depth was approximately 0.2 m, and approximately 600 g soil was collected for per sample. Before and after our experiment, samples were collected and sealed in polyethylene packages, and then transported to the laboratory as soon as possible to analyze the soil total nitrogen, phosphorus, potassium, and the concentration of Cd. The first samples were collected at different test plots in June after three days of remediation, and the second samples were collected in September when the rice paddies had been harvested (100 days after the first samples). The first/second samples from each site were pooled, homogenized into triplicate samples, and respectively placed in sterile 50 ml tubes and stored at −20°C until further molecular analyses and physicochemical property analyses could be performed.

The parameters involved in this experiment were obtained with several methods. The concentration of Cd (total and available) was determined by ICP-OES (IRIS INTREPID II XDL THERMO) following the method described in EPA 3052 (15). The pH value was measured in an aqueous extract (soil/water=1/5 w/v).The soil total nitrogen (TN) was determined by the Kjedahl method (16). Soil total phosphorus (TP) concentration was determined by alkaline digestion followed by molybdate colorimetric measurement (17). The soil total potassium (TK) was determined using atomic absorption spectrophotometry (18). As for the soil available phosphorus (AP), it was first extracted with 0.5 M NaHCO_3_ (19) and then determined by molybdenum colorimetric method (20). Available nitrogen (AN) was determined following extraction in 2 M KCl (21), while available potassium (AK) was determined by flame photometry (22). The soil organic matter (OM) content was determined by K dichromate oxidation method (23).

### RNA extraction and metagenomic analysis

Total community genomic RNA extraction was performed using a E.Z.N.A.Soil RNA Kit (Omega, USA), following the manufacturer’s instructions. We measured the concentration of the RNA using a Qubit 2.0 (Life,USA), to ensure that adequate amounts of high-quality genomic RNA had been extracted. The V3-V4 region of the 16S rRNA gene was amplified using using KAPA HiFi Hot Start Ready Mix (2×) (TaKaRa Bio Inc., Japan). AMPure XP beads were used to purify the free primers and primer dimer species in the amplicon product and then samples were delivered for library construction using universal Illumina adaptor and index. Before sequencing, the RNA concentration of each PCR product was determined using a Qubit^®^ 2.0 Green double-stranded RNA assay and it was quality controlled using a bioanalyzer (Agilent 2100, USA). Sequencing was performed using the Illumina MiSeq system (Illumina MiSeq, USA), according to the manufacturer’s instructions. The sequences obtained in this study were deposited in the GenBank database under accession number PRJNA396262. For taxonomic analysis, the 16S rRNA gene sequences of each sample would be submitted to the RDP Classifier to assign archaeal and bacterial taxonomy to 97%. Shannon index was calculated using mothur. For functional analysis, KEGG (Kyoto Encyclopedia of Genes and Genomes) database were used to assign joined reads to different metabolic pathways in PICRUSt, and only the annotated sequences were used for further analysis (24).

### Statistical analyses

Principal coordinates analysis (PCoA) and the unweighted pair group method with arithmetic mean (UPGMA) were performed by R (v.2.13.1; http://www.r-project.org/) with packages VEGAN to explore the taxonomic and functional relationships between the samples (25). Redundancy analysis (RDA) was conducted by Canoco5.0 to identify the key environmental variables shaping the taxonomic compositions. An analysis of variance (ANOVA) was used to calculate the significance of Shannon index and Welch’s t-test within VEGAN was used to calculate the differences in relative abundance of genes and species between samples, and p < 0.05 was considered to be statistically significant. F-statistic, indicating the variance among soils to variance within a field, was calculated by analysis of variance (ANOVA) models.

## RESULTS

### Soil Cd Content Change Before and After Amendment

The distribution of total cadmium (TCd) and available cadmium (ACd) in our experimented soil (Fig. 2; Table S1) showed that the average T_Cd_ content before the experiment ranged from 0.96 (field E) to 1.11 (field C) mg/kg, which was significantly higher than 0.3 mg/kg set by the secondary standard of Environmental Quality Standards for Soils (EQSS) (26). Moreover, the highest content of A_Cd_ among the groups was group C, with the average content of 0.7 mg/kg, while the lowest was in the control group (CK1) with 0.54 g/kg. After the experiment, the T_Cd_ content was consistent with that of the initial, but the ACd content was significantly decreased. Among them, amendment A performed best with a significant decrease of 41.27%, followed by B, D, and C, with decreases of 36.67%, 34.55%, 32.86%, respectively, while amendment E performed worst, with a decrease of merely 29.42%. Meanwhile, the contents of T_Cd_ and A_Cd_ in CK2 decreased simultaneously, but no significant changes were observed in CK1. Thus, it was possible that ACd in the CK2 group was absorbed by rice. Besides, the coefficients of variation (CVs) of pH, TN, TP, and TK in our experimental soils (Table S1) were all less than four, indicating that the soil was relatively homogeneous.

**FIG 2.**
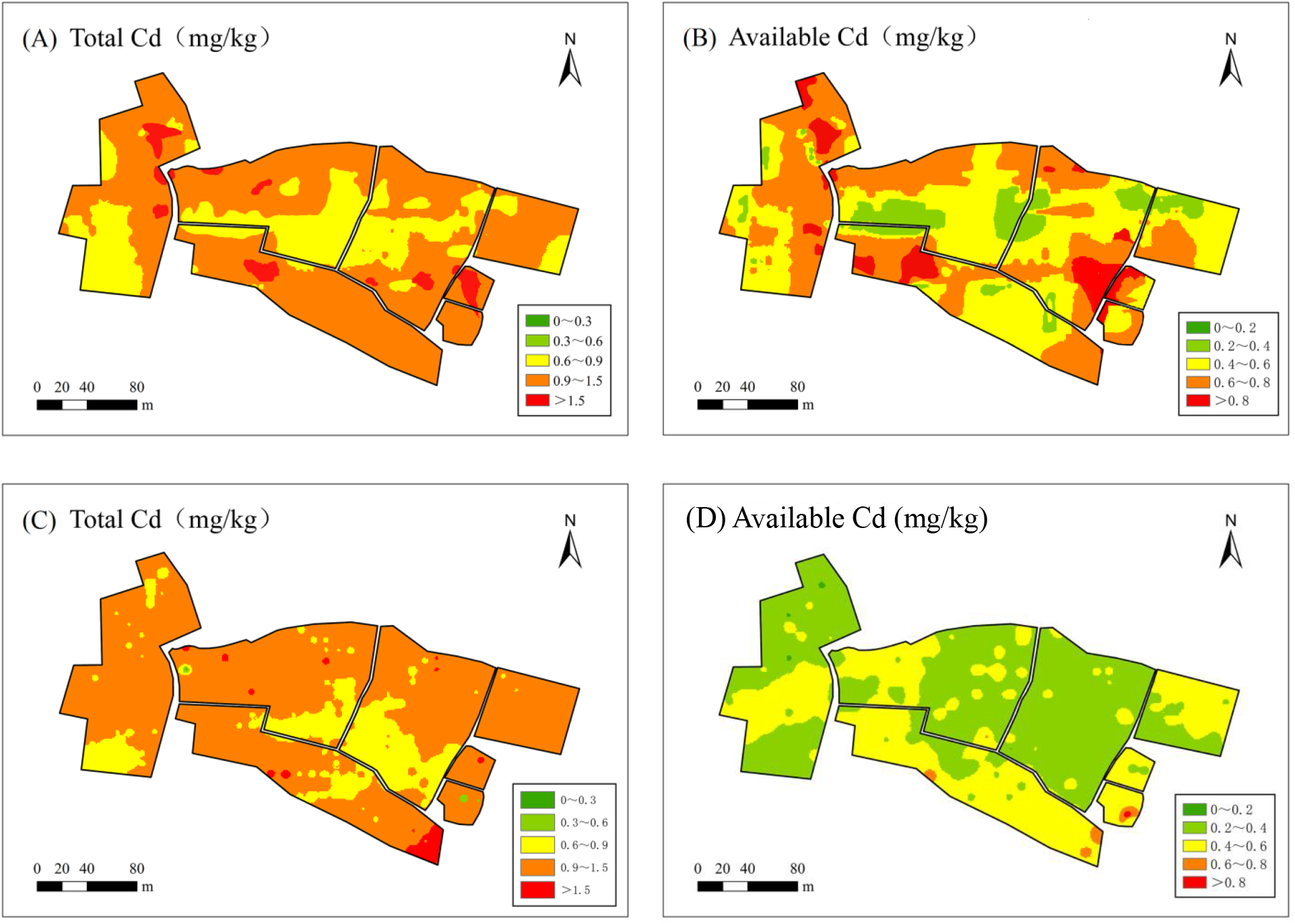
The distribution of total cadmium and available cadmium in our experimented soil. (A) and (B) showed the distribution of cadmium in the 1^st^ samples, (C) and (D) showed the distribution of cadmium in the 2^nd^ samples.

### Physicochemical Properties of the Collected Samples

The soil physicochemical properties in the first and second samples were analyzed (Table 2). In addition to CK1, the soil TN in the first samples showed the greatest change among all measured variables with a CV1 of 25.20, while the soil OM varied the least with a CV1 of 4.86, indicating the greatest impact of the amendments was on soil TN. Specifically, field A had the highest TN level and pH value. Field D had the lowest TN, TK, TP, AN, AP, and OM contents. Field E had the lowest soil pH value.

**Table 2.** Soil physicochemical properties of samples from each sampling site.

| Sample | The first samples (g/Kg) |  |  |  |  |  |  |  | The second samples (g/Kg) |  |  |  |  |  |  |  |
| --- | --- | --- | --- | --- | --- | --- | --- | --- | --- | --- | --- | --- | --- | --- | --- | --- |
|  | pH | TN | TK | TP | AK | AN | AP | OM | pH | TN | TK | TP | AK | AN | AP | OM |
| A1 | 7.67 | 2.13 | 18.85 | 1.52 | 0.15 | 0.26 | 0.05 | 42.16 | 7.24 | 2.27 | 17.20 | 2.38 | 0.13 | 0.18 | 0.04 | 42.16 |
| A2 | 7.24 | 2.10 | 17.79 | 1.49 | 0.14 | 0.25 | 0.05 | 41.20 | 6.83 | 2.14 | 16.23 | 2.25 | 0.12 | 0.17 | 0.03 | 41.20 |
| A3 | 7.44 | 2.08 | 18.26 | 1.47 | 0.15 | 0.26 | 0.05 | 42.66 | 7.02 | 2.20 | 16.67 | 2.31 | 0.12 | 0.18 | 0.04 | 42.66 |
| B1 | 7.43 | 1.88 | 17.82 | 1.76 | 0.17 | 0.29 | 0.05 | 42.61 | 7.43 | 2.27 | 17.10 | 2.43 | 0.13 | 0.18 | 0.04 | 42.61 |
| B2 | 7.01 | 1.90 | 16.82 | 1.78 | 0.16 | 0.27 | 0.05 | 43.08 | 7.01 | 2.14 | 16.14 | 2.29 | 0.13 | 0.17 | 0.04 | 41.68 |
| B3 | 7.20 | 1.89 | 17.27 | 1.88 | 0.16 | 0.28 | 0.05 | 42.06 | 7.20 | 2.20 | 16.57 | 2.35 | 0.13 | 0.18 | 0.04 | 42.66 |
| C1 | 6.53 | 1.99 | 19.36 | 1.62 | 0.16 | 0.27 | 0.05 | 46.33 | 6.56 | 2.29 | 17.20 | 2.41 | 0.13 | 0.19 | 0.04 | 45.33 |
| C2 | 6.16 | 1.93 | 18.27 | 1.52 | 0.15 | 0.25 | 0.05 | 47.40 | 6.19 | 2.16 | 16.23 | 2.28 | 0.12 | 0.18 | 0.04 | 46.80 |
| C3 | 6.33 | 1.88 | 18.76 | 1.67 | 0.16 | 0.26 | 0.05 | 47.02 | 6.36 | 2.22 | 16.67 | 2.34 | 0.13 | 0.18 | 0.04 | 46.32 |
| D1 | 7.54 | 1.23 | 17.41 | 1.59 | 0.18 | 0.23 | 0.05 | 41.33 | 7.14 | 2.23 | 16.89 | 2.37 | 0.13 | 0.18 | 0.04 | 41.33 |
| D2 | 7.12 | 1.18 | 16.43 | 1.58 | 0.17 | 0.22 | 0.04 | 41.19 | 6.74 | 2.11 | 15.94 | 2.24 | 0.12 | 0.17 | 0.04 | 40.19 |
| D3 | 7.31 | 1.20 | 16.87 | 1.60 | 0.17 | 0.23 | 0.05 | 40.77 | 6.92 | 2.16 | 16.37 | 2.30 | 0.13 | 0.18 | 0.04 | 41.17 |
| E1 | 6.29 | 1.42 | 17.61 | 1.66 | 0.18 | 0.25 | 0.05 | 42.47 | 6.84 | 2.25 | 16.79 | 2.38 | 0.13 | 0.18 | 0.04 | 42.47 |
| E2 | 5.94 | 1.34 | 16.62 | 1.69 | 0.17 | 0.23 | 0.05 | 42.36 | 6.45 | 2.12 | 15.84 | 2.25 | 0.12 | 0.17 | 0.04 | 42.06 |
| E3 | 6.10 | 1.36 | 17.07 | 1.70 | 0.17 | 0.24 | 0.05 | 41.82 | 6.63 | 2.18 | 16.27 | 2.31 | 0.12 | 0.18 | 0.04 | 43.52 |
| CK1-1 | 7.30 | 1.03 | 17.20 | 1.43 | 0.18 | 0.24 | 0.05 | 41.40 | 7.47 | 2.15 | 16.38 | 1.32 | 0.15 | 0.16 | 0.05 | 41.31 |
| CK1-2 | 6.89 | 1.14 | 16.23 | 1.40 | 0.17 | 0.23 | 0.05 | 41.24 | 7.05 | 2.03 | 15.45 | 1.25 | 0.14 | 0.15 | 0.05 | 40.27 |
| CK1-3 | 7.08 | 1.11 | 16.67 | 1.59 | 0.17 | 0.23 | 0.05 | 40.88 | 7.24 | 2.08 | 15.87 | 1.28 | 0.15 | 0.16 | 0.05 | 41.84 |
| CK2-1 | 6.93 | 1.18 | 16.79 | 1.43 | 0.18 | 0.22 | 0.04 | 42.59 | 6.32 | 2.16 | 17.30 | 2.51 | 0.11 | 0.14 | 0.03 | 41.88 |
| CK2-2 | 6.54 | 1.02 | 15.84 | 1.54 | 0.17 | 0.21 | 0.04 | 40.67 | 5.97 | 2.04 | 16.33 | 2.36 | 0.10 | 0.14 | 0.02 | 40.97 |
| CK2-3 | 6.72 | 1.15 | 16.27 | 1.40 | 0.17 | 0.22 | 0.04 | 39.95 | 6.13 | 2.09 | 16.77 | 2.43 | 0.10 | 0.14 | 0.03 | 40.82 |
| CV | 7.47 | 26.84 | 5.46 | 8.26 | 6.36 | 8.97 | 5.32 | 4.81 | 6.42 | 3.40 | 3.03 | 17.65 | 10.43 | 8.84 | 18.08 | 4.26 |
| CV1 | 7.96 | 25.20 | 5.57 | 7.89 | 6.61 | 7.99 | 5.67 | 4.86 | 6.25 | 3.12 | 2.68 | 3.09 | 8.05 | 8.79 | 13.94 | 4.42 |

CV (coefficient of variation) is used to calculate the variations between all samples, higher value represents greater variation.
CV1 indicates the variations between samples except the control CK1. Abbreviations: TN, total nitrogen; TP, total phosphorus; TK,
total potassium; AN, available nitrogen; AP, available phosphorus; AK, available potassium; OM, organic materials.

However, the variations in soil pH, TN, TK, TP, and OM collected in the second samples were smaller than those in the first samples, indicated by their CV1 value. The CV1 of soil TN had the biggest change, reduced from 25.20 to 2.68, followed by the CV1s of soil pH, TK, TP and OM, indicating that the effect of amendments on those factors decreased significantly after three months. Beyond that, the CV1s of the first samples were similar to those of CV, whereas the CV1s of the second samples, especially that of TP, did not coincide with CV. This indicated that variations among the first samples resulted mainly from amendments, but the impact of the amendments was minimized after three months, and planting rice acted as an influential factor affecting the physicochemical parameters of the soil.

### Microbial Community Diversity

In our experiment, the total number of full-length quality sequences obtained from the 42 samples was 3,335,373 in bacteria and 1,725,544 in archaea, with ranges of 28,139-136,334 and 21,104 and 64,729 sequences, respectively. After rigorous filtering of low-quality reads, barcode and primer trimming, and filtering of the chimeras, a total of 3,180,967 sequences of bacteria (average read length, 420bp) and 15,874,876 sequences of archaea (average read length, 380bp) were grouped and classified into operational taxonomic units (OTUs) at the 97% similarity level. The Shannon index (Fig. 3) responding to variables resulting from experimental procedures of amendment revealed that the archaeal and bacterial Shannon index in field E showed a significant decrease compared to that in the control CK2, while the Shannon indices in other fields were slightly improved. About three months later, a more similar diversity level was maintained in second samples, except for in field E, which still had a significantly low bacterial diversity (P < 0.05). This indicates that the impact of all amendments on microbial diversity could be weak and varied in bacteria and archaea over time.

**FIG 3.**
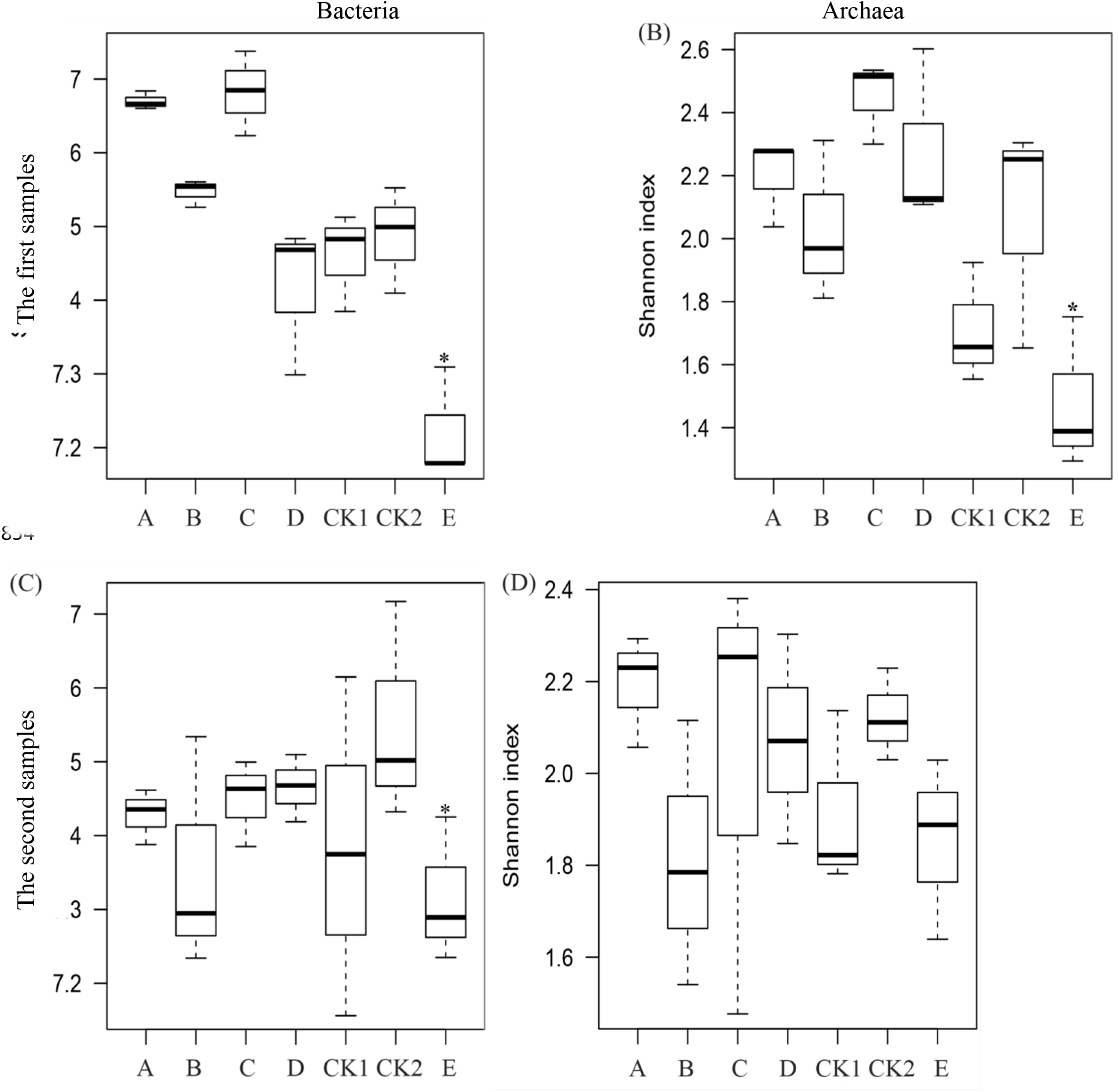
The bacterial and archaeal Shannon index of all samples collected at different times. * indicates index had significant variation compared with CK2 tested by ANOVA (P<0.05)

### Composition and taxonomic features of bacteria and archaea

The most abundant bacterial organisms in all samples (Fig. 2A; Table S3) were from *Proteobacteria* phylum, accounting for 37.73%-47.40% of the total valid reads in all libraries. Specifically, *Proteobacteria* from the first samples showed significantly higher abundances in amended soils compared to that in the control CK2, especially in field E (46.36%), while there was no marked difference between amended soils and CK2 from the second samples. Within the *Proteobacteria, Alphaproteobacteria* from the first samples (Fig. S1) showed relatively higher abundances in B, D, and E, compared to that in the control CK2, while *Alphaproteobacteria, Deltaproteobacteria,* and *Gammaproteobacteria* from the second samples showed relatively higher abundances in D, A, and E compared to that in the control CK2. It can be seen that there were significant differences between classes of soil *Proteobacteria* in different soil amendments. Frequently, variations in the relative abundance of other phyla compared to those in CK2 were also observed. For example, the relative abundance of *Gemmatimonadetes* in all amended soils from the first samples declined significantly, but less variation was observed in the second samples. Other phyla, such as *Acidobacteria, Firmicutes, Chloroflexi, Verrucomicrobia,* and *Nitrospirae,* responded differently to different soil amendments. In addition to bacteria (Fig. 4B; Table S4), *Thaumarchaeota* was the most abundant phylum in all samples, accounting for 68.6 to 96.16% of the total valid reads in the first samples but 56.33 to 74.25% in the second samples, and *Thaumarchaeota* from field E showed the highest abundance in both the first samples and second samples. *Crenarchaeota* also was detected in all samples and was less abundant in field in both the first samples and second samples compared to in the control.

**FIG 4.**
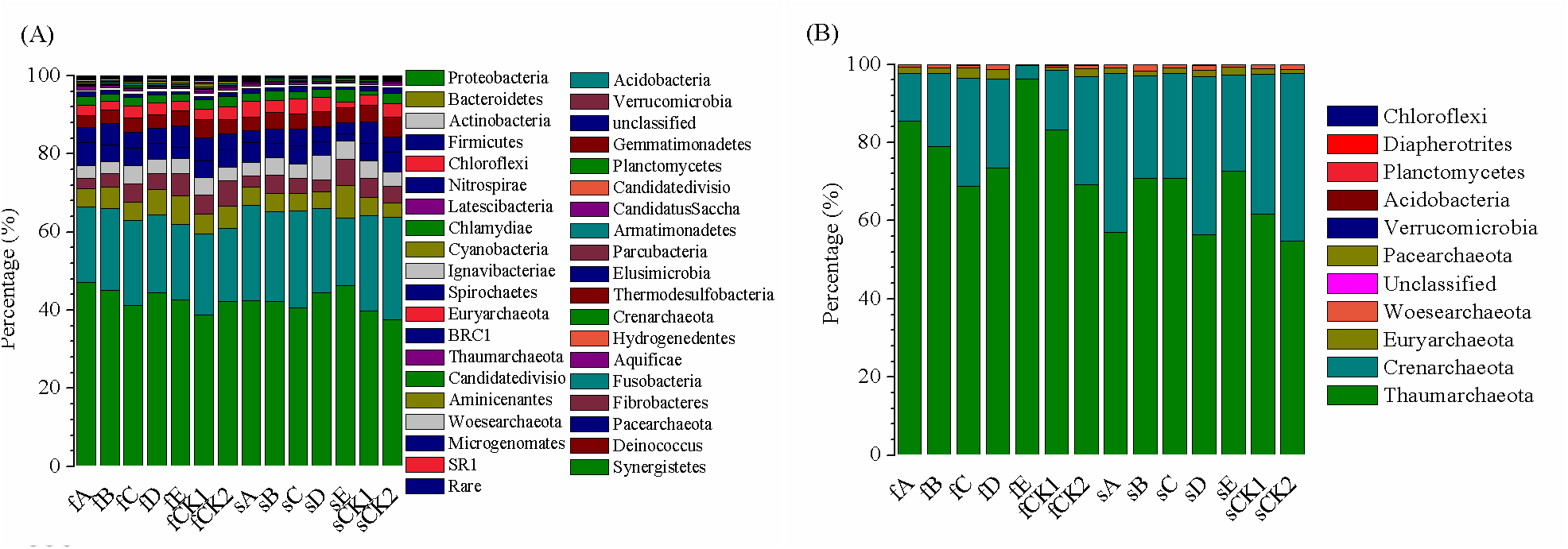
Taxonomic classification of mean bacterial (A) and archaeal (B) reads retrieved from different samples at phylum level. The prefix was used to distinguish samples collected at different periods with the first letter of the “first” and “second”. And the percentage is an average of the three sets of data. “Rare” indicates phyla with relative abundances of less than 0.01%.

At the genus level, the microbial compositions also differed significantly across the amended soils and the control (Fig. S1; see also Table S5 and S6 in the supplemental material). Among the genera identified, *Gp6* showed a relatively high abundance (> 3%) in all bacterial samples, especially in the library retrieved from fA (> 8.4%). *Sphingomonas* and *Gemmatimonas,* which have relative abundances over 2% in all of the sequencing libraries, saw significant increase and decrease in the first samples, respectively, especially in fE (8.31% and 3.78%). *Nitrososphaera* was the most abundant archaeal genus (> 55%) in all samples, especially in the libraries retrieved from fE (96.09%) and sE (74.13%). Other genera only demonstrated obvious changes in relatively high abundances in part of samples. For example, the lowest abundances of the sequences of *Gp16* were found in libraries from fE (0.88%).

### Functional classification

Through computerized processing of known biological processes in the cell and standardizing the explanation of existing gene functions, we systematically analyzed the functions of the gene and classified the gene functions into different levels with the BIOENV procedure within the R package vegan. At level 2 (Fig. 5; Table S7 and Table S8), most of the active genes both in archaea and bacteria were associated with metabolism, followed by genetic information processing and environmental information processing, while cellular processes, human diseases, and organismal systems took approximately 12.5% (bacteria) and 5.9% (archaea) of the total. As shown in Fig. 4, the most active microbial metabolisms were mainly amino acid metabolism, carbohydrate metabolism and energy metabolism both under the stable flooded and dry states. Moreover, the gene expression of energy metabolism in field D and E were significantly lower than that in the control CK2.

**FIG 5.**
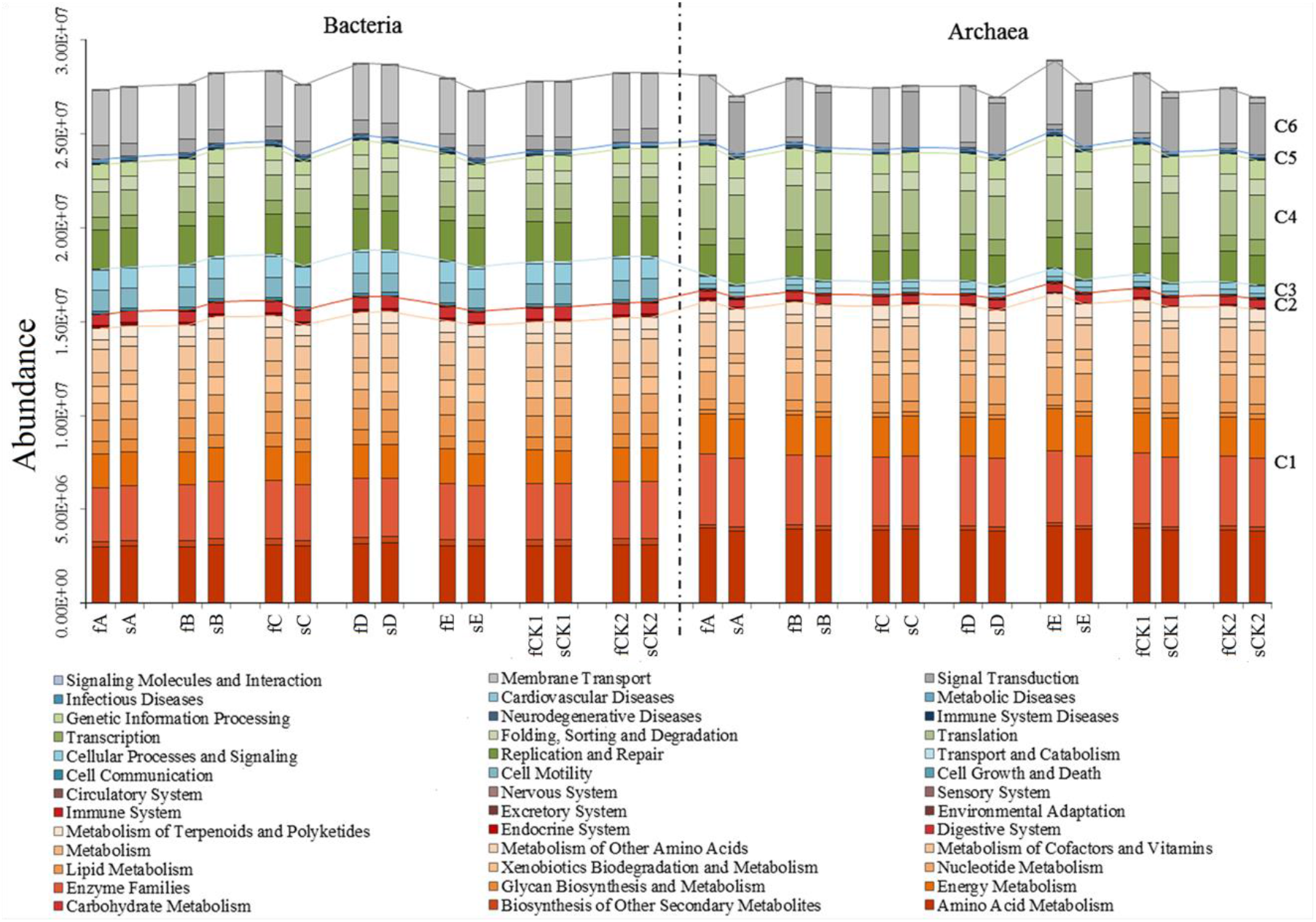
Gene functional categories of microorganism in experimented paddy fields referring to KEGG level 2. C1 to C6 represent 6 broad gene functional categories distinguished by different color, which were Metabolism, Organismal Systems, Cellular Processes, Genetic Information Processing, Human Diseases and Environmental Information Processing respectively. Samples collected in June and September were represented symbolically by first letter of the “first” and “second”.

The genes of carbohydrate, energy, and amino acid metabolisms in our amended soils at level 3 were further analyzed, and the results (Table S9 and S10) showed that there were a total of 10 pathways for energy metabolism, 16 pathways for carbohydrate metabolism, and 24 pathways for amino acid metabolism. The most abundant genes of bacteria and archeae in all samples were related to oxidative phosphorylation (> 1.6%) and methane metabolism (> 2.1%), respectively, both of which belonged to the energy metabolism pathway. Carbon fixation in photosynthetic organisms was also another pathway with relative abundances over 1% in all samples. The most abundant genes of protein metabolism and carbohydrate metabolism were related to amino acid-related enzymes (> 1.2%) and amino sugar and nucleotide sugar metabolism (> 1.1%), respectively. The difference between the amended soils and CK2 was determined by Welch’s t-test, which indicated that the number of pathways of archaea with significant differences was less than that of bacteria, and the number of pathways with significant differences from the first samples was much greater than that from the second samples. For the pathways of the first bacterial samples, the most distinct one from CK2 was E, followed by B, A, and D, and the difference between field C and CK2 was the smallest. E and D showed a significant decrease in nitrogen metabolism, methane metabolism, oxidative phosphorylation, and carbon fixation pathways in prokaryotes compared to those in CK2, while the gene expression of metabolic pathways in other amended soils were lower or only slightly higher than those in CK2.

### Variation of taxonomic and functional patterns and their correlations with environmental variables

Principal coordinate analysis (PCoA) based on Bray-Curtis distance was used to visualize the distances and variations between samples (Fig. 6), and samples collected in June and September were represented symbolically by first letter of the “first” and “second”. The comparison of the microbial communities among the 42 bacterial samples indicated that amendments played important roles in the shaping of microbial communities in the paddy soils. The field E harbored communities, clustered clearly apart from the others, while the fields A and C were clustered close to the controls CK1 and CK2. In addition, the first bacterial samples (Fig. 6A) showed a sparser distribution than that of the second samples, indicating that three days of remediation by amendment could significantly stimulate the activity of soil microorganisms, while after three months, the physicochemical environment of the soil and the whole soil ecosystem have time to stabilize, which might lead to a weaker response to amendments. In addition to bacteria, though most archaeal samples (Fig. 6B) were clustered disorderly and irregularly, samples from field E had the least similarity of microbial communities compared to others, and also indicating that the bacteria can respond more quickly and significantly to the changes in soil environment in amended paddy fields.

**FIG 6.**
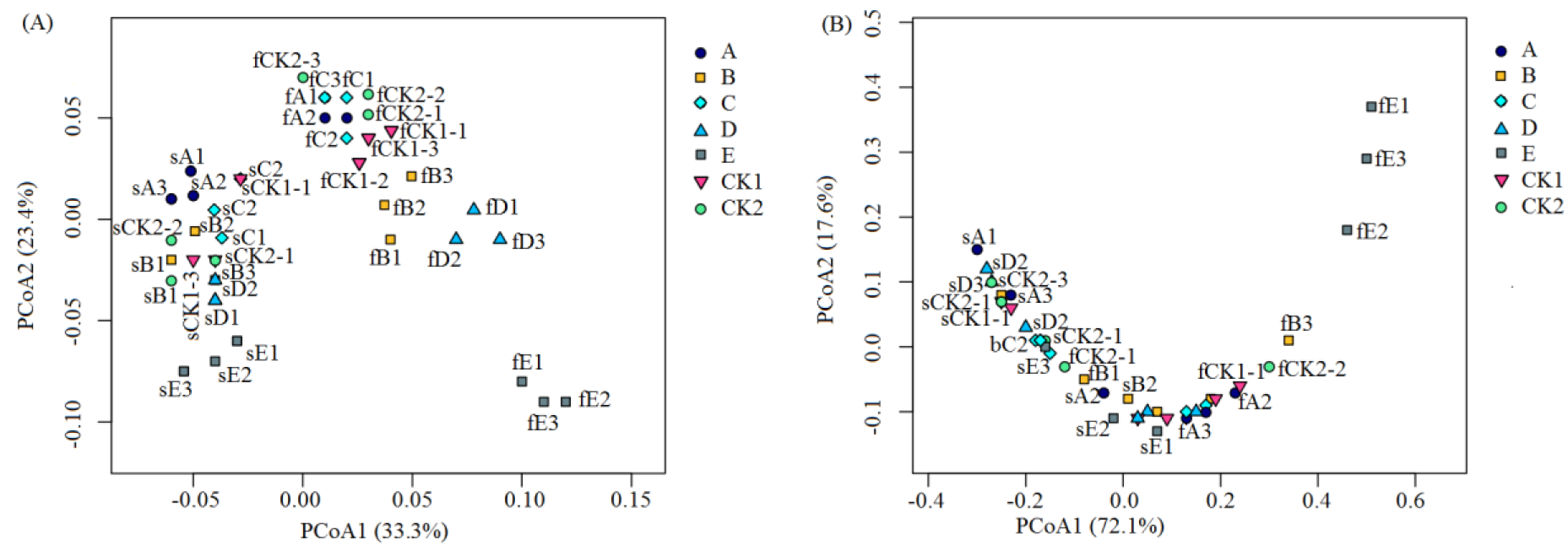
The Principal Co-ordinates Analysis (PCoA) of bacteria (A) and archaea (B) based on unifrac dictance (3D). Different colors of spot represent samples from different fields. The more gathered in the figure, the higher the similarity between samples, otherwise, similarity between samples would be lower when the space distance is far. Samples collected in June and September were represented symbolically by first letter of the “first” and “second”.

Differences between 42 functional genes were demonstrated from the 16S rRNA species composition data. The UPGMA tree (Fig. S2) indicated a similar result from PCoA. When examining the first bacteria UPGMA tree, samples from fA, fC and the two control groups (fCK1 and fCK2) clustered, and samples from fB and fD clustered. However, the microbial function of the fE group was distinct from others. When rice paddies were harvested, the longest distance of the sample of sE and the shortest distance of the sample of sCK1 and sC in bacteria implied that the amendments in E could have a sustainable influence on the bacterial gene function. Samples in the archaea UPGMA tree were clustered disorderly with a shorter branch compared to that of bacteria, illustrating that the least number of variables occurs in archaea. However, the fE group was distinct from its counterparts and the control groups, indicating that the greatest impact on microbial function within a short time was caused by amendment E.

The above-mentioned dating results implied that, generally, functional patterns had similar results to taxonomic patterns by taking all the samples into consideration. Therefore, one could predict the overall functional patterns from the taxonomic composition of the microbiomes, and similar conclusions were drawn from previous studies analyzing other ecosystems such as biogas processes and soil and gut microbiomes (29,30,31). Thus, redundancy analysis (RDA) was used to determine the environmental variables that most shaped the taxonomic patterns (Fig. 7). Dominant bacterial and archaeal phyla were chosen from Table S3 and S4 on the basis of relative abundance > 1%. The RDA model explained 82.8% and 83.4% of the total variance in taxonomic patterns of bacteria and archaea, respectively. Of the eight selected environmental variables, pH, by a relatively large magnitude, appears to be the most important environmental variable for both archaeal and bacterial taxonomic patterns. In addition, the AK, TP, and TN also respectively explained more than 10% of the total variance in both archeae and bacteria. *Acidobacteria, Proteobacteria,* and *Bacteroidetes* were positively correlated with higher TN and negatively correlated with pH, which may suggest that acidic amendments may facilitate the growth of these bacteria. However, *Chloroflexi, Verricomicrobia, Nitrospirea, Woesearchaeota,* and *Euryarchata* were positively correlated with pH, indicating that these bacteria favor alkaline environments.

**FIG 7.**
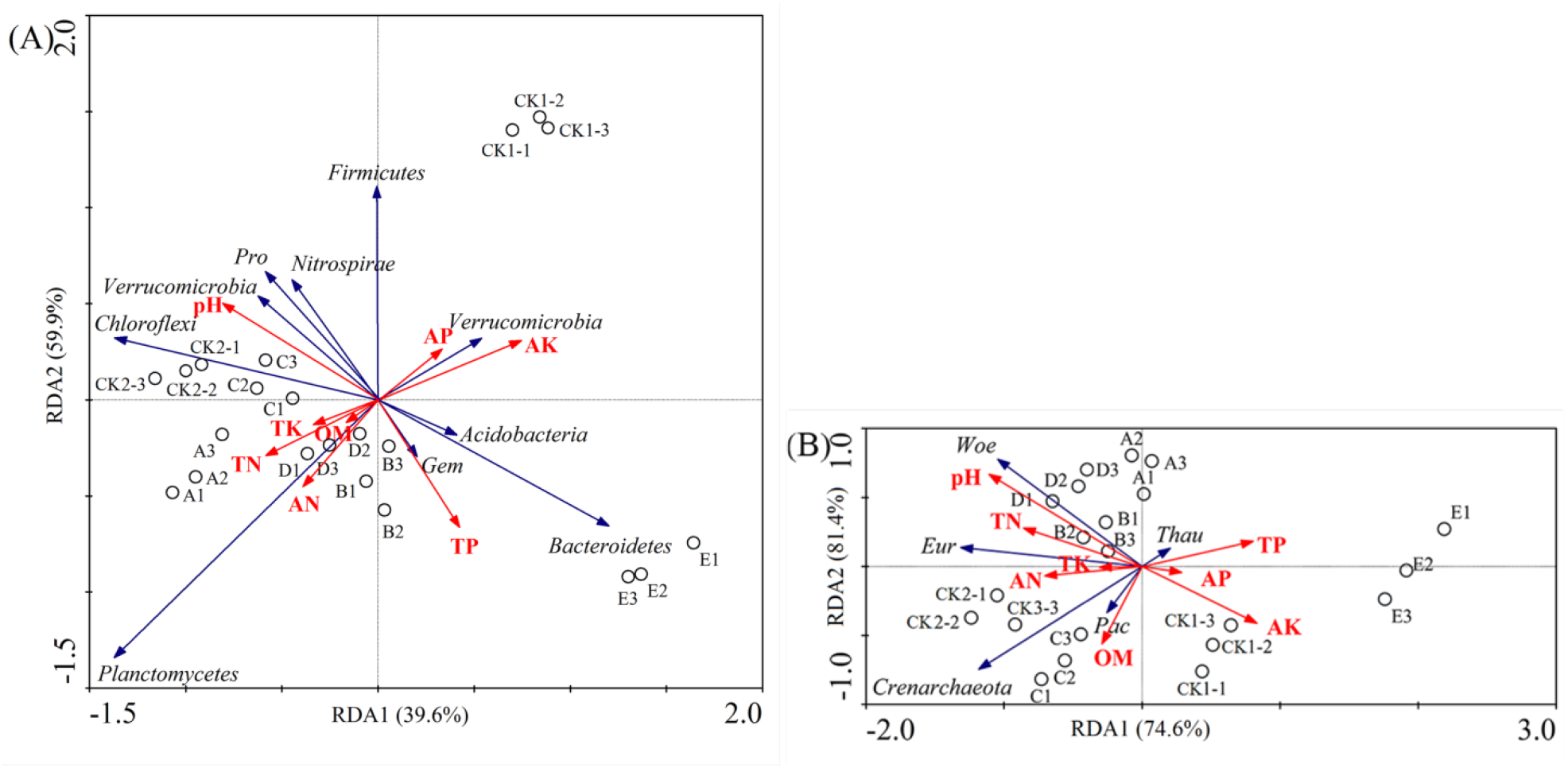
Ordination diagrams from redundancy analysis (RDA) of bacterial (A) and archaeal (B) relative abundance and physicochemical parameters. Red vectors represent the magnitude of parameters while black vectors represent communities. Each group is represented by black circles. (A) RDA analysis of bacteria with Axis 1 and 2 explained 39.6% and 59.9% of total variation respectively. (B) RDA analysis of archaea with Axis 1 and 2 explained 74.6% and 81.4% of total variation respectively. Abbreviations: *Pro, Proteobacteria*; *Gem, Gemmatimonadetes*; *Thau, Thaumarchaeota*; Eur, *Euryarchaeota*; *Woe, Woesearchaeota; Pac, Pacearchaeota.*

Our ANOVA analysis of diversity-environmental factors (Table S2) showed that the soil pH was the main reason for the change in soil bacterial and archaeal diversity, which was reflected by PCCs (Pearson correlation coefficients) of 0.8899 in bacteria and 0.5213 in archaea, with a significant difference (P value <0.01) in our experiment. Additionally, others were positively correlated with diversity, except for TP and TK. However, three months later, none of the factors had a significant impact on biodiversity, indicating that the influence of environmental variables caused by amendments on microbial diversity had decreased.

## DISCUSSION

### Environmental data that shape the microbial community composition

At present, due to the sharp contradictions in population, food, resources, and environment, the sustainable development of agriculture has received unprecedented attention, and the soil quality has become a major issue facing all mankind. As an important part of soil ecological systems, soil microorganisms can respond to soil changes caused by amendments sensitively and rapidly, and the relationship between them and soil metabolism will contribute to agricultural development priorities in the future. In this paper, the impacts of five typical soil amendments on microbes during the process of remediation were studied in the field. In fact, similar studies have been reported in cadmium-contaminated soils, which proved that microbial diversity could be higher than expected when using soil amendments consisting of red mud, zeolite, and lime (32). Those studies also proved that using brown amphibolite and iron sand in Cd-contaminated soil at lower levels had a weak influence on bacterial abundance (33), implying that the effects on microflora structure and diversity varied sharply among soil physicochemical properties determined by amendments. In our experiment, amendment A most significantly improved soil microbial diversity, followed by C and B, while E showed an opposite effect. Our ANOVA analysis of diversity-environmental factors showed that soil pH was the main reason for the change in soil bacterial and archaeal diversity, which is in accordance with the result from Griffiths (34) on factors affecting the repair of Cd-contaminated rice fields by soil amendment.

Due to their narrow tolerance ranges to soil pH, bacterial community structures will be greatly and easily affected changes in pH (35). In a study on the distributional orderliness of 14 typical geographic soils at different altitudes in eastern China using the phospholipid fatty acid analysis method, Wu (36) found that differences in soil microbial community structures are mainly caused by soil pH, and gram-negative bacteria (GNB) increased the pH. Jones (37) discovered that pH is the most important environmental factor affecting the composition of *Acidobacteria*, which is one of the most advantageous soil bacterium groups globally, contributing 20% of the 16s rRNA clone library of bacteria on average, and can even reach 80% in some soils. In this study, pH explained 14.9% of soil bacterial changes and 17.1% of archaea changes on the level of the phylum, which makes it the most remarkable factor affecting soil bacteria and archaea communities.

Additionally, RDA analysis indicated that, together, AK, TN, and TP explained 34.7% of variance in bacteria and 39.0% in archaea at the phylum level, particularly those of *Acidobacteria, Proteobacteria,* and *Woesearchaeota.* The contents of soil N, P, and K are closely related to microbial community structure. Zhong et al. (38) found that the average well color development value and Shannon index under P-deficient conditions were significantly lower than normal. Chen et al. (39) also found that significant changes occurred in microbial community structure and function in red paddy soil, reflected by the lower number and activity of bacteria under P-deficient conditions than those under normal conditions. In this experiment, A, B, and C amendments applied nutrient material, such as phosphates, while amendments D and E contained certain minerals to provide more niche for soil microbes, especially for *Proteobacteria* (40). On the other hand, an obvious difference in AP content in different amended soils was observed due to the phosphate material in amendment, and the increase in soil AP meets the phosphorus requirement of plant growth; thus, in the plant, the metabolism would get stronger, be accompanied by a large amount of photosynthetic activity, and sent nutrients to the root system to promote the physiological activities related to root growth and increase the rhizosphere effect. It is supposed that secretion into the rhizosphere can promote the growth of microorganisms, thereby promoting the increase of *Proteobacteria* (41). Interestingly, well-developed root systems also increased the crop’s demand for potassium and nitrogen, which can lead to a relative reduction in soil AK and AN; this could be one reason for the lower AK content in A, B, and C amended soils than in the control group and D and E amended soils.

### Dominant genes functions in paddy field

The analysis of the main microbial metabolisms in paddy soil showed that the amino acid metabolism, carbohydrate metabolism and energy metabolism were the core of the metabolic activity under both the stable flooded and dry states. Similarly, these dominant metabolisms were also observed in the microbiomes of other environments such as biogas plants (42), activated sludge in wastewater treatment system (43), desert soil (44) and deep-sea sediments (45), which are supposed to be comparable with flooded paddy soil in terms of anaerobic conditions. Other metabolisms most closely related to the three core metabolisms, and the assimilation and metabolism of soil nutrients were potassium, methane, sulfur, nitrogen, and phosphorus, which all showed a high expressional level for amino acid, carbohydrate, and energy metabolisms, which was attributed to the fact that nitrogen and sulfur are the nutrients mainly involved in protein biosynthesis; phosphorus is a key component (adenosine triphosphate, ATP) of energy metabolism; and potassium plays an important role in maintaining the osmotic balance in cells (46).

### Biogeochemical N-cycling in amended soil

In our study, the change in TN in the soil of each experimental group in this study was significant, indicating dynamic biogeochemical N cycling in amended soils. Soil nitrogen (47), as one of the most essential elements in the crop growth process and the most deficient nutrient element in soil, directly affects the growth of crops in terms of its abundance and supply. Additionally, previous research has reported that organic nitrogen can combine with heavy metals to form a stable compound with a high affinity, such as Cu^2+^, in soil (48). Although microorganisms have not been isolated in this study, it may be possible to understand the effect of amendments based on functional genes and closely related microorganisms. The genes involved in nitrogen metabolism were barely expressed in archaea (Fig. 8), indicating that bacteria contributed more to nitrogen metabolism.

**FIG 8.**
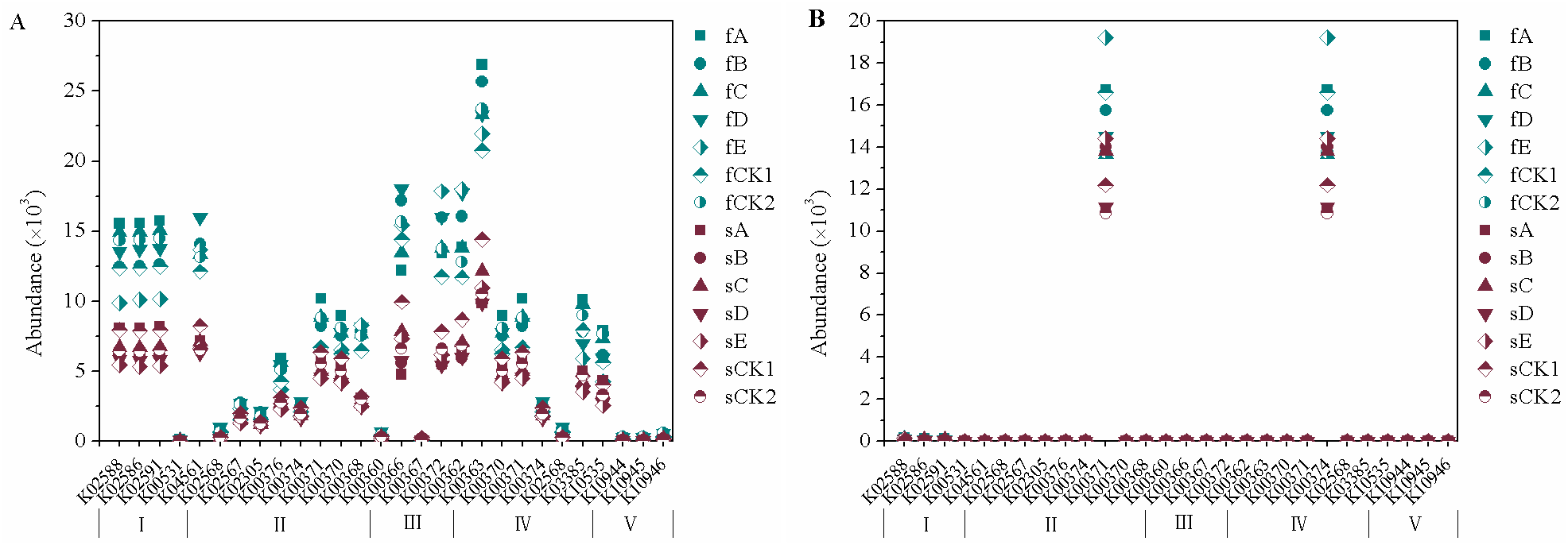
Genes involved in soil N-cycling pathways from metagenomic datasets of all bacterial (A) and archaeal (B) samples. The abundance of genes is average abundance. I, II, III, IV, and V respectively represent pathway of Nitrogen fixation, Denitrification, Assinilatory nitrate reduction, Dissimilatory nitrate reduction and Nitrification. Samples collected in June and September were represented symbolically by first letter of the “first” and “second”.

### Nitrogen fixation

Research has shown that rhizobia can secrete protons and various organic acids, amino acids, which contribute to dissolving heavy metals by increasing the acidity of the soil, or change the form of heavy metals by coordination with metabolites (49). In our study, in addition to the higher expression of individual genes in the nitrogen reduction, the expression of genes related to nitrogen fixation in each experimental group was higher than that of the rest. This is possibly because a large part of nitrogen intake by rice is derived from nitrogen fixation by nitrogen-fixing microorganisms (50). Soil nitrogen fixation is an important source of soil AN and an important part of soil N cycling (51). Different from leguminous plants, the main biological nitrogen fixation in paddy fields is associated with nitrogen fixation, which mainly depends on the nitrogen-fixing microorganisms in the soil (52, 53). Therefore, the variations in species and structure of nitrogen-fixing microorganisms in soil have an important influence on N cycling in rice fields (54). Nitrogen-fixing bacteria, such as *Bradyrhizobium*, *Rhizobium*, *Mesorhizobium*, *Geobacter*, *Clostridium* and *Anaeromyxobacter* were detected as most the abundant genera in this experiment (Table S11), and most of these bacteria belonging to *Alphaproteobacteria.* This is consistent with the results of Rösch et al (55). Specifically, though *Bradyrhizobium, Rhizobium* showed relatively higher abundances in amended field E, the accumulative abundance of nitrogen-fixing microorganisms was higher in several libraries from the soil samples taken from the amended fields A and B, which may be related to the pH value of the amendments. Bannert et al. (56) recently explored the differentiation of nitrogen-fixing bacteria communities and their quantities in the tidal flat wetlands and paddy fields of Zhejiang Province in China, finding that microbial communities and quantities were largely correlated with pH.

### Nitrogen reduction

Because of the unique tillage management system, rice paddy fields are under flooded conditions most of the time, with intermittent drying, and long dry and wet alternation causes redox reactions in rice soil. At the same time, the effect of secreted oxygen on the rice rhizosphere also places the paddy soil in a state of aerobic/anaerobic co-existence for a long time, so that severe redox reactions of nitrogen elements and the nitrate reduction process can cause the production of greenhouse gases N^2^ and N^2^O and cause a series of problems such as nitrogen loss and an increase in greenhouse gas production. According to the different microbial physiological processes, nitrate reduction, includes assimilatory nitrate reduction, dissimilatory nitrate reduction and denitrification. In this study, the gene expression of denitrification in all samples was relatively low, and soil nitrate reduction was given priority to with dissimilatory nitrate reduction (DNRA). Studies have shown that assimilatory nitrate reduction is extremely sensitive to NH_4_^+^; a soil NH_4_^+^ concentration of 0.1 mg/kg can inhibit the assimilatory nitrate reduction by 60% within 1 min, while 10 mg/kg inhibits it by 80% after 5 min (57). Flooded soils usually contain rich amounts of NH_4_^+^ and organic nitrogen, so nitrate assimilation reduction is unlikely to be the dominant one in the soil at the ecological level, which is consistent with our results. At the same time, the community of NRB showed similar results. The dominant NRB in all samples were *Thiobacillus*, *Pseudomonas*, and *Bacillus*, which were the commonly detected DNRA bacteria genera (58, 59). Compared to the results in CK2, *Pseudomonas* showed significantly higher abundances in several samples taken from the first samples and amended soil sA, but *Pseudomonas* was less abundant in both fE and sE.

Within the DNRA, nirD (K00363, Nitrite reductase (NADH), EC:1.7.1.15) was the most abundant gene in all samples, of which A had the highest gene expression, indicating that soil microbes in A amended soil can be motivated to adapt to sudden anaerobic stress accompanied by changes the soil in the redox status; in this case, the denitrifying enzyme systems were activated quickly, using a large amount of NADH accumulated during aerobic respiration to ensure the denitrification process for anaerobic respiration (60). Moreover, the weakest adaptability in the E field may be related to low soil pH. For example, Qin et al. found that the acidification of soil could lead to decreased activity in NADH (61). Bergaust et al. pointed out that the NADH of Paracoccus denitrificans was difficult to assemble at lower pH levels (62), resulting in a low enzyme activity. Other alternative explanations may relate to low-density and porous materials in amendment E, which can optimize aeration conditions in the soil, lead to a lower anaerobic degree, and thereby inhibit the NADH activity (63).

### Nitrification

Soil nitrification is an important part of the nitrogen cycle, and has a great influence on the environment. Soil NO_3_^−^ produced in the process of nitrification is leached more easily than NH_4_^+^, and causes serious loss of nitrogen in the soil. On the other hand, a high level of nitrification can reduce the soil pH value, leading to the release of heavy metals and reducing the effect of amendments. In our study, the most abundant nitrobacteria in all samples phylogenetically belonged to the *Nitrospira* and *Nitrosospira*, but the abundance of nitrobacteria is far lower than that of nitrogen-fixing bacteria. Specifically, the relative abundance of *Nitrosospira* in fields D and E were significantly lower than that in control group CK2. At the same time, the gene expressions of nitrification in all samples were lower than those of other metabolites. *Hao* (K10535, hydroxylamine dehydrogenase, EC: 1.7.2.6), as the gene with the highest expression, was observed to have the lowest abundance in E. Prior studies have shown that pH strongly influences the metabolism of nitrifying bacteria (64). Soil nitrification is not obvious in soils with a pH value from4.6-5.1, and is relatively slow in soils with a pH value from 5.8-6.0; however, it is strong in soils with a pH value from6.4-8.0 (65). Additionally, different pH values of soils contribute to the differences in activity and species of nitrifying bacteria. In soils with lower pH values, the number and activity of autotrophic nitrifying bacteria are lower, resulting in weak nitrification. It is generally believed that the optimal pH for autotrophic nitrifying bacteria growth ranges from 6.6-8.0 (66).

In conclusion, due to fact that the rice field is an entire ecosystem, applying amendments has an impact on the structure and function of the soil ecosystem, from the energy flow to the material cycle, and thus could indirectly affect soil microbes; for example, this could make the nitrogen-fixing bacteria in the soil more capable of nitrogen fixation than those in conventional (non-amended) rice fields, and the utilization ratio of soil nitrogen can be increased.

### Environmental effects of amendments

Metagenomic technology, as a method of exploring microbial community structures based on the gene diversity of microbes, has become a global research hotspot in recent years. Compared with traditional pure culture techniques, metagenomic technology helps in understanding the interaction relationship between microbial and polluted environment, and generating information that indicates the degree of pollution in the environment, to monitor and evaluate the environment. A growing number of studies (67) confirm that climate change will affect biological species, and the response of a single species in the isolated state to environmental change will be differentiated from those of species in an interaction relationship. Only by further studying the relationships between environmental stress factors and the composition of microbial communities at the genome level can we truly understand complex microbial populations and their complex relationships with biology, the environment, and ecology, and also understand their spatial distribution and dynamic change, which will provide theory for environmental monitoring, warning, and assessment.

In our study, alkaline amendments (A and B) composed of CaO had significant positive effects on the decrease of soil Acd, which can be probably due to pH value. A large number of studies (68, 69) have shown that soil pH is a very important factor affecting available heavy metal concentration, controlling the adsorption-desorption of heavy metals in soil and the chemical behaviors such as dissolution-deposition. And alkaline amendments also had a favorable impact on taxonomic and functional patterns, especially on microbial diversity, soil nitrogen fixation and the growth of related bacteria, which conducive to the decrease of soil Acd. However, amendment A, largely composed of calcite (CaCO^3^), performed better than amendment B, indicating that the environmental effect was different due to their different composition proportions. Application of calcite can improve the soil pH and increase the surface negative charge of soil particles, prompt the Cd in the soil to transform into hydroxide or carbonate precipitation, which decreases the bioavailability of Cd and ultimately improves the microbial characteristics of the soil (70). However, phosphorous materials have been reported to be used in acidic soil; thus, pH limited the effect of amendment B on largely phosphorous material.

Different alkaline amendments may have different effects. Alkaline amendment D, based on MgO (magnesium oxide), showed a lower removal rate of soil A_cd_ and inhibited the gene expression of soil nitrogen fixation, nitrification, and the growth of related bacteria. A possible reason for this may be that soil would be hardened, wet, muddy, and crusting with poor permeability during the application of the magnesia alkalinity amendment (71, 72), which could be harmful to microbial growth and their matter and energy exchange with the environment (73, 74).

The differences in taxonomic and metabolic patterns in group C and the control group CK2 were minimal, with little difference in the individual functional classification, while that in group E varied significantly, which could have been caused by the minimum difference in the average soil pH, and smaller or insignificant differences in AN and TN between C and CK2. Amendment E, with the lowest pH value, was the main cause of reductions in the microbial diversity, nitrogen fixation, nitrogen reduction, nitrification, and the growth of related microorganisms.

At present, the remediation of soil heavy metals by amendments has been further researched, its technologies are becoming increasingly developed, and the remediation effect has also increased gradually. However, the problem with seeking perfection is that amendments can lead to other problems in the soil environment. In the practical application, the pH value, the type of amendment, and the proportion of components must be taken into consideration in order to achieve the most suitable effect both ecologically and effectively.

### Conclusion

This study reports the variation in taxonomic and functional patterns of microbiomes by metagenomic sequencing across CCPFs restored with five amended soils, revealing that pH is the major factor affecting the microbial communities examined in this study and in other studies (36, 37). Furthermore, a number of N-metabolizing bacteria exhibited high abundance in amended soils, and DNRA saw a great positive response to nitrogen and pH from amendments, which is in accordance with the current understanding of the metabolic capacities of indigenous microorganism. Importantly, the response of microbial communities to amendments was weak after three months, which is critical to elucidating the roles of active taxa in the functions of amended ecosystems and optimizing the management of soil amendments. After the remediation for 3 months, the soil physiochemical environment and soil ecosystem began to stabilize because of soil buffering or the downward migration of amendment ingredients, occurring when amended soils were turned over during rainfall or irrigation, consequently the response of microbial communities to amendments was weak. And unmeasured factors like hydrothermal conditions, rhizosphere effects, and crop growth may have a more temporal influence on the structure and diversity of different microbial communities (27, 28). However, atmospheric oxygen can diffuse steadily into flooded paddy soil surfaces at least 2 mm deep, forming a tiny aerobic zone (75, 76). Because of time constraints and limitations of the technology, this study failed to distinguish the microbial process of aerobes and anaerobes when soil was flooded, and merely mixed topsoil and deep soil homogeneously instead, to study the microbial community of aerobes and anaerobes. In the future, we need to conduct further studies on the main physiological metabolic processes in flooded CCPFs.

## ACKNOWLEDGMENTS

This work was supported by the National Natural Science Foundation of China (No. 41271094), Soil Environmental Protection Engineering Technology Center in Sichuan. Particular thanks the EPA in Mianzhu and their villagers for allowing us sample their fields and assisting with management and date access. Thanks also to Sangon Biotech (Shanghai) Co.,Ltd for field assistance of Illumina MiSeq sequencing technology.

## REFERENCES

1. Suthar S, Nema AK, Chabukdhara M, Gupta SK. 2009. Assessment of metals in water and sediment of Hindon River, India: Impact of industrial and urban discharges. Journal of Hazardous Materials 171:1088–1095, https://doi.org/10.1016/jjhazmat.2009.06.109

2. Vollmann J, Lošák T, Pachner M, Watanabe D, Musilová L, Hlušek J. 2015. Soybean cadmium concentration: validation of a QTL affecting seed cadmium accumulation for improved food safety. Euphytica 203(1): 177–184. https://doi.org/10.1007/s10681-014-1297-8

3. Nicholson FA, Smith SR, Alloway B.J, Smith CC, Chambers BJ. 2003. An inventory of heavy metals inputs to agricultural soils in England and Wales. The Science of the Total Environment 3:205–219. https://doi.org/10.1016/S0048-9697(03)00139-6

4. Cao HC, Luan ZQ, Wang JD, Zhang XL. 2009. Potential ecological risk of cadmium, lead and arsenic in agricultural black soil in Jilin Province, China. Stochastic Environmental Research and Risk Assessment 23(1):57–64. https://doi.org/10.1007/s00477-007-0195-1

5. Lin AJ, Zhang XH, Su YH, Hu Y, Cao Q, ZhuYG. 2007. Chemical fixation of metals in soil using bone char and assessment of the soil genotoxicity. Environmental Science 28:232–237. http://www.ncbi.nlm.nih.gov/pubmed/17489175

6. Zhu YG, Chen BD. 2000. Bioremidiation Technologies of Contaminated Soils. Beijing: Environmental Science Press of China 2000:11–42.

7. Zhang L, Li L, Pan G. 2009. Variation of Cd, Zn and Se contents of polished rice and the potential health risk for subsistence-diet farmers from typical areas of South China, China, Environ. Sci 30:2792–2797. (in Chinese)

8. Hasegawa H, Rahman IM M, Rahman MA. 2015. Environmental Remediation Technologies for Metal-Contaminated Soils. Environmental Remediation Technologies for Metal-Contaminated Soils (pp. 1–254). Springer Japan. https://doi.org/10.1007/978-4-431-55759-3

9. Gu JG, Zhou QX, Wang X. 2003. Reused path of heavy metal pollution in soils and its research advance. Journal of Basic Science and Engineering 11:143–151.

10. Long XX, Yang XE, Ni WZ. 2002. Current situation and prospect on the remediation of soils contaminated by heavy metals. Chinese of Applied Ecology 13:757–762. (in Chinese)

11. Wang L, Xu YM, Sun GH, Liang X, Sun Y. 2012. Immobilization of cadmium and lead compound contaminated soil using new porous material combined with phosphate. In 2012 2nd International Conference on Remote Sensing, Environment and Transportation Engineering. https://doi.org/10.1109/RSETE.2012.6260594

12. Zhao RF, Zou CQ, Zhang FS. 2007. Effects of long-term P fertilization on P and Zn availability in winter wheat rhizoshpere and their nutrition. Plant Nutrition and Fertilizer Science 13(3):368–372. (in Chinese)

13. Huang CM, He LR, Wen AB. 1993. The classification of the degradation of purple soil in Sichuan. Mountain Research 11:201–208. (in chinese)

14. SPMS. Sichuan Public Meteorology Service. 2016. Available at: http://pcc.scqx.gov.cn/sc/shouye

15. Gil C, Boluda R, Ramos J. 2004. Determination and evaluation of cadmium lead and nickel in greenhouse soils of Almería (Spain). Chemosphere 55:1027–1034. https://doi.org/10.1016/_j.chemosphere.2004.01.013

16. Bremner, J. M. 1996. Nitrogen-total. In: Sparks, D.L. (Ed.), Methods of Soil Analysis. Part 3. Chemical Methods. Soil Science Society of America, American Society of Agronomy, Madison, WI, 1085–1121.

17. Murphy J, Riley JP. 1962. A modified single solution method for the determination of phosphate in natural water. Analytica Chimica Acta 27:31–36.

18. Helmke P, Sparks D. 1996. Methods of soil analysis. Part 3. Chemical methods, issue 5. Published by SSSA and ASA. 551–574.

19. Olsen SR, Cole C, Watanabe FS, Alan LA. 1954. Estimation of available phosphorus in soils by extraction with sodium bicarbonate. USDA Circular 939.

20. Kuo S. 1996. Phosphorus. In: Sparks, D.L. (Ed.), Methods of Soil Analysis. Part 3. Chemical Methods. Soil Science Society of America, American Society of Agronomy, Madison, WI, 869–919.

21. Page AL, Miller RH, Keeney DR. 1982 Methods of Soil Analysis. Part 2. Chemical and Microbiological Properties. American Society of Agronomy, Soil Science Society of America, Madison, WI.

22. Knudsen D, Peterson GA, Pratt PF. 1982. Lithium, sodium and potassium. Methods of Analysis 2:225–246.

23. Yeomans JC, Bremner JM. 1988. A rapid and precise method forroutine determination of organic carbon in soil. Communications in Soil Science and Plant Analysis 19:1467–1476. https://doi.org/10.1080/00103628809368027

24. Fierer N, Leff JW, Adams BJ Nielsen UN, Bates ST, Lauber CL. 2012. Cross-biome metagenomic analyses of soil microbial communities and their functional attributes. Proc Nat AcadSci USA 109:21390–21395. https://doi.org/10.1073/pnas.1215210110

25. Kuczynski J, Stombaugh J, Walters WA, González A, Caporaso JG, Knight R. 2012. Using QIIME to analyze 16S rRNA gene sequences from microbial communities. Curr Protoc Microbiol 27:1E.5.1–1E.5.20. http://dx.doi.org/10.1002/9780471729259.mc01e05s27.

26. EQSS. 1995. Environmental Quality Standards for soils. Minister of Environmental Protection of the People’s Republic of China. Available at: http://english.sepa.gov.cn/Resources/standards/Soil (in Chinese).

27. Liu SH, Xie PF, Xu GH, Wang RL. 2013. The effect of pH on the growth characteristics of seedling roots of hybrid rice seedlings. Jiangsu Agricultural Sciences. 41:70–72.

28. Ross DJ, Speir TW, Kettles HA, Tate KR, Mackay AD. 1995. Soil microbial biomass, C and N mineralization, and enzyme-activities in a hill pasture - Influence of grazing management. Soil Biology & Biochemistry. 27:1431–1443.

29. Muegge BD, Kuczynski J, Knights D, Clemente JC, González A, Fontana L, et al. 2011. Diet drives convergence in gut microbiome functions across mammalian phylogeny and within humans. Science 332(6032):970–974.

30. Fierer N, Lauber CL, Ramirez KS, Zaneveld J, Bradford MA, Knight R. 2012. Comparative metagenomic, phylogenetic and physiological analyses of soilmicrobial communities across nitrogen gradients. ISME J 6:1007–1017.

31. Gilbert JA, Field D, Swift P, Thomas S, Cummings D, Temperton B, et al. 2010. The taxonomic and functional diversity of microbes at a temperate coastal site: a ‘multi-omic’ study of seasonal and diel temporal variation. PLoS ONE 5(11):e15545.

32. Gnmu G, Castaldl P, Sontona L, Deiana P, Melis P. 2007. Influence of redmud, zeolite and lime on heavy metals immobilization culturable heterotrophic microbial population and enzyme activities in a contaminated soil. Geoderma 142:47–57. https://doi.org/10.1016/j.geoderma.2007.07.011

33. Mench M, Renella G, Gelsomino A, Landi L, Nannipieri P. 2006. Biochemical parameters and bacterial species richness in soils contaminated by sludge-borne metals and remediated with inorganic soft amendments. Environmental Pollution 144, 24–31. https://doi.org/10.1016/j.envpol.2006.01.014

34. Griffiths R, Thomson B, James P, Bell T, Bailey M, Whiteley A. 2011. The bacterial biogeography of British soils. Environmental microbiology 13: 1642–1653. https://doi.org/10.1111/j.1462-2920.2011.02480.x

35. Ahn JH, Song J, Kim BY, Kim MS, Joa JH, Weon HY. 2012. Characterization of the bacterial and archaeal communities in rice field soils subjected to long-term fertilization practices. Journal of Microbiology. 50:754–765. https://doi.org/10.1007/s12275-012-2409-6

36. Wu Y, Ma B, Zhou L, Wang H, Xu J, Kemmitt S, Brookes PC. 2009. Changes in the soil microbial community structure with latitude in eastern China, based on phospholipid fatty acid analysis. Applied Soil Ecology. 43:234–240. https://doi.org/10.1016/j.apsoil.2009.08.002

37. Jones RT, Robeson MS, Lauber CL, Hamady M, Knight R, Fierer N. 2009. A comprehensive survey of soil acidobacterial diversity using pyrosequencing and clone library analyses. ISME Journal 3:442–453. https://doi.org/10.1038/ismej.2008.127

38. Zhong W, Cai Z. 2007. Long-term effects of inorganic fertilizers on microbial biomass and community functional diversity in a paddy soil derived from quaternary red clay. Applied Soil Ecology 36: 84–91. https://doi.org/10.1016/j.apsoil.2006.12.001

39. Chen JG, Zhang YZ, Zeng XB, Zhou WJ, Tan ZJ, Jiang DS. 2008. Effects of different fertilizations on soil microbial characteristics in a paddy soil from red earth with long-term K-deficiency. Plant Nutrition and Fertilizer Science 14 (6):1200–1205. (in Chinese)

40. Rosello MR, Aman, R. 2001. The species concept for prokaryotes. FEMS Microbial Reviews 25: 39–67. https://doi.org/10.1016/S0168-6445(00)00040-1

41. Zhao YH, Ling QH. 2011. Response of Abundance and Composition of the Bacterial Community to Long-term Fertilization in Paddy Soils. Scientia Agriculture Sinica 44: 4610–4617. (in Chinese)

42. Luo G, FotidisI A, Angelidaki I. 2016. Comparative analysis of taxonomic, functional, and metabolic patterns of microbiomes from 14 full-scale biogas reactors by metagenomic sequencing and radioisotopic analysis. Biotechnology for Biofuels. 9:1–12.

43. Ju F, Guo F, Ye L, Xia Y, Zhang T. 2014. Metagenomic analysis on seasonal microbial variations of activated sludge from a full-scale wastewater treatment plant over 4 years. Environ Microbiol Rep 6:80–90.

44. Fierer N, Leff JW, Adams BJ, Nielsen UN, Bates ST, Lauber CL, Owens S, Gilbert JA, Wall DH, and Caporaso, JG. 2012. Crossbiome metagenomic analyses of soil microbial communities and their functional attributes. Proc Nat AcadSci USA 109:21390–5. https://doi.org/10.1073/pnas.1215210110

45. Lesniewski RA, Jain S, Anantharaman K, Schloss PD, Dick GJ. 2012. The metatranscriptome of a deep-sea hydrothermal plume is dominated by water column methanotrophs and lithotrophs. The ISME Journal 6: 2257–2268. https://doi.org/10.1038/ismej.2012.63

46. Kamil MAL, Jobori, Salim SEA. 2014. The effect of Water limitation on Water Relations, growth and seed yield of four soybean (Glycine max merri.) genotypes. Advances in Bioresearch 88:371–380.

47. Bueno E, Mesa S, Bedmar E. J. Richardson DJ, Delgado MJ. 2012. Bacterial adaptation of respiration from oxic to microoxic and anoxic conditions: redox control. Antioxidants & Redox Signaling 16(8):819–852. https://doi.org/10.1089/ars.2011.4051

48. Wang B, Zhao J, Guo Z, Ma J, Xu H, Jia Z. 2015. Differential contributions of ammonia oxidizers and nitrite oxidizers to nitrification in four paddy soils. ISME Journal 9(5): 1062–1075. https://doi.org/10.1038/ismej.2014.194

49. Guo XJ, Huang QY, Zhao ZA, W Chen. 2001. Effect of microorganisms on the mobility of heavy metals in soil environments. China J Appl Environ Biol 8(1): 105–110.

50. Bodelier PLE. 2003. Interaction between oxygen-releasing roots and microbial processes in flooded soils and sediments. Ecological Studies, Root Ecology 168:331–362.

51. Segata N, Izard J, Waldron L, Gevers D, Miropolsky L, Garrett W S, Huttenhower C. 2011. Metagenomic biomarker discovery and explanation. Genome Biology. 12(6). https://doi.org/10.1186/gb-2011-12-6-r60.

52. Jin J, Tai JC, Pan GX, Li LQ, Song XY, Xie T, Liu XY, Wang D. 2012. Comparison of Soil Organic Carbon, Microbial Diversity and Enzyme Activity of Wetlands and Rice Paddies in Jingjiang Area of Hubei, China. Scientia Agricultura Sinica 45 (18): 3773–3781.

53. Han B, Kong JJ, Zou XM, Gong HD. 2009. The Evolvement and Expectation of Biological Nitrogen Fixation. Journal of Shanxi Agricultural Sciences 37(10): 86–89.

54. Yuan M, An SJ, Sun JG. 2016. Isolation and Biological Properties of Endophytic Diazotrophs from Rice and Their Influences on Rice Seedling Cd Accumulation. Scientia Agricultura Sinica 49 (19): 3754–3768.

55. Rösch C, Mergel A, Bothe H. 2002. Biodiversity of denitrifying and dinitrogen-fixing bacteria in an acid forest soil. Applied and Environmental Microbiology 68 (8): 3818–3829. https://doi.org/10.1128/AEM.68.8.3818-3829.2002

56. Bannert A, Kleineidam K, Wissing L, Mueller-Niggemann C, Vogelsang V, Welzl G, Cao Z, and Schloter M. 2011. Changes in diversity and functional gene abundances of microbial communities involved in nitrogen fixation, nitrification, and denitrification in a tidal wetland versus paddy soils cultivated for different time periods. Applied and Environmental Microbiology 77(17): 6109–6116. https://doi.org/10.1128/AEM.01751-10.

57. Rice CW, Tiedije JM. 1989. Regulations of nitrate assimilation by ammonium in soils and in isolated soil microorganisms. Soil Biology and Biochemistry 21:579–602. https://doi.org/10.1016/0038-0717(89)90135-1.

58. Kelly DP, Wood AP. 2000. Confirmation of Thiobacillus denitrificans as a species of the genus Thiobacillus, in the beta-subclass of the Proteobacteria, with strain NCIMB 9548 as the type strain. International Journal of Systematic and Evolutionary Microbiology 50:547–550. https://doi.org/10.1099/00207713-50-2-547.

59. Silver WL, Herman DJ, Firestone MK, Ecology S, Sep N. 2012. Dissimilatory Nitrate Reduction to Ammonium in Upland Tropical Forest Soils DISSIMILATORY NITRATE REDUCTION TO AMMONIUM IN UPLAND TROPICAL FOREST SOILS. Ecology 82(3): 2410–2416. https://doi.org/10.2307/2679925

60. Hesterberg D, Chou JW, Hutchison KJ, Sayers DE. 2001. Bonding of Hg(2) to reduced organic, sulfur in humic acid as affected by S/Hg ratio. Environmental science and Technology 35(13): 2741–2745. https://doi.org/10.1021/es001960o

61. Karlsson T, Elgh-Dalgren K, Björn E, Skyllberg U. 2007. Complexation of cadmium to sulfur and oxygen functional groups in an organic soil. Geochimica Et Cosmochimica Acta 71:604–614. https://doi.org/10.1016/j.gca.2006.10.011

62. Wang J, Li Y, Liang C. 2008. Recovery of transgenic plants by pollen-mediated transformation in Brassica juncea. Transgenic Research 17(3):417–424. https://doi.org/10.1007/s11248-007-9115-x

63. Laird D A, Fleming P, Davis D D, Horton R, Wang B, Karlen D L. 2010. Impact of biochar amendment on the quality of a typical Midwestern agricultural soil. Geoderma 158: 443–449. https://doi.org/10.1016/j.geoderma.2010.05.013.

64. Koops HP, Purkhold U, Pommerening-Röser A, Timmermann G, Wagner M. 2006. The Lithoautotrophic Ammonia-Oxidizing Bacteria. In The Prokaryotes. New York, NY: Springer New York. pp. 778–811. https://doi.org/10.1007/0-387-30745-1_36

65. Katyal C, Cater MF, Vlek PLG. 1988. Nitrification activity in submerged soils and its relation to denitrification loss. Biology and Fertility of Soils 7(1): 16–22. https://doi.org/10.1007/BF00260726.

66. Li LM, Pan YH. 2000. Nitrification and influencing factors in Taihu Lake. Soils 19:289–293. (in chinese)

67. Treusch AH, Kletzin A, Raddatz G, Ochsenreiter T, Quaiser A, Meurer G, Schleper C. 2004. Characterization of large-insert DNA libraries from soil for environmental genomic studies of Archaea. Environmental Microbiology 6(9): 970–980. https://doi.org/10.1111/j.1462-2920.2004.00663.x

68. Ahn JY, Kang SH, Hwang KY, Kim HS, Kim JG, Song H, Hwang I. 2015. Evaluation of phosphate fertilizers and red mud in reducing plant availability of Cd, Pb, and Zn in mine tailings. Environmental Earth Sciences 74(3):2659–2668. https://doi.org/10.1007/s12665-015-4286-x

69. Liang Z, Peng X, Luan Z. 2012. Immobilization of Cd, Zn and Pb in sewage sludge using red mud. Environmental Earth Sciences 66(5):1321–1328. https://doi.org/10.1007/s12665-011-1341-0

70. Wang CW, Xu YM, W L. 2010. Effect of immobilization contaminated soil by cadmium and lead using sepiolite and phosphate. Journal of Safety and Environment 10:42–45.

71. Hinrich L, Bohn BL, Mcneal et al. 2001. Soil chemistry. John wiley & Sons, inc. 35–40

72. Zhang XC, Norton LD. 2002. Effect of exchangeable Mg on saturated hydraulic conducitivity, disaggregation and clay dispersion of disturbed soils. Journal of Hydrology 260:194–205. https://doi.org/10.1016/S0022-1694(01)00612-6.

73. Marcar NE, Termaat A. 1990. Efects of root zone solutes on Eucalyptus camaldulensis and Eucalyptus bicostata seedlings:Response to Na^+^,Mg^2+^ and Cl^−^. Plant soil 125:245–254. https://doi.org/10.1007/BF00010663v.

74. Pankhurst CE, Yu S, Hawke BG, Harch BD. 2001. Capacity of fatty acid profiles and substrate utilization patterns to describe differences in soil microbial communities associated with increased salinity or alkalinity at three locations in South Australia. Biology and Fertility of Soils 33:204–217 https://doi.org/10.1007/s003740000309.

75. Bomberg M, Jurgens G, Saano A, Sen R, Timonen S. 2003. Nested PCR detection of Archaea in defined compartments of pine mycorrhizospheres developed in boreal forest humus microcosms. FEMS Microbiology Ecology 765 43:163–171. https://doi.org/10.1016/S0168-6496(02)00384-7

76. Chelius MK, Triplett EW. 2001. The diversity of Archae and bacteria in association with the roots of Zea mays L. Microbial Ecology 41:252–263. https://doi.org/10.1007/s002480000087

